# Scalable Non-negative Matrix Factorization of the Human Cell Census Reveals Interpretable Transcriptional Programs

**DOI:** 10.64898/2025.12.30.697085

**Authors:** Yu-Ting Liu, Timothy J Triche, Zachary J DeBruine

## Abstract

Large single-cell atlases now span tens of millions of cells, yet few provide reusable and interpretable reference representations that support direct biological reasoning at atlas-scale. Here, we present an interpretable Non-negative Matrix Factorization reference embedding of 28.5 million healthy cells and approximately 60,000 genes from the Human Cell Census. The resulting gene and cell weights define additive transcriptional programs that align with annotated cell types and organized biological pathways. New datasets can be projected into this fixed reference space without fine-tuning or retraining, as we demonstrate using an independent cystic fibrosis dataset. This resource provides a transparent coordinate system for exploratory analysis and hypothesis generation, complementing deep embeddings that prioritize integration or prediction with a representation designed for interpretability and reuse.

## Introduction

Single-cell RNA sequencing has enabled cell-resolved analysis of gene expression across tissues, developmental stages, and physiological states. Large coordinated efforts have consolidated these data into atlas-scale resources, including the Human Cell Atlas and the Human Cell Census as accessed through the cellxgene infrastructure [1, 2]. These atlases support comparative analysis across biological contexts but increasingly expose a gap between data availability and the availability of modeling representations that allow atlas-scale structure to be interrogated, summarized, and reused in a transparent way [3, 4].

Latent representations are central to most analyses of large single-cell collections. Deep generative models provide flexible embeddings that support denoising, integration, and probabilistic inference across heterogeneous datasets [5, 6]. Foundation-model approaches further emphasize transferability across large collections of single-cell profiles [7, 8, 9]. These models are generally optimized for expressivity and predictive performance, but their latent dimensions are not typically designed to correspond to stable, human-interpretable biological programs [10, 11, 12]. Our objective here is not batch correction or task-specific prediction, but a reusable reference space with interpretable factors.

A useful way to distinguish these goals is to view a latent representation as an interface between raw measurements and downstream reasoning. Many modern embeddings prioritize expressivity—capturing nonlinear variation to support integration or prediction within the framework of a specific model architecture—but do not guarantee that individual dimensions correspond to reusable biological objects or insights. In contrast, a reference embedding is most valuable when its coordinates are stable and human-interpretable, and even model-agnostic: a factor can be named, inspected, aggregated, and reused as a consistent unit of analysis across studies [13, 14]. This paper focuses on that complementary objective, treating factor identity as an explicit design constraint rather than an emergent byproduct of model flexibility.

Non-negative matrix factorization (NMF) produces an interpretable decomposition in which non-negativity induces learning of sparse gene programs that reconstitute transcriptional state when added together [15, 16, 17, 18]. Early biological applications demonstrated that NMF-derived metagenes capture well-organized molecular patterns [19], and subsequent work in single-cell analysis has shown that such factors can separate cell identity from activity and support transfer across platforms, tissues, and species [16]. In this setting, interpretability is operational: gene weights define ranked transcriptional programs that can be annotated directly, and cell weights define compositional summaries that can be aggregated across annotations without additional modeling assumptions [13, 20].

The need for such representations is amplified by the scale and heterogeneity of contemporary atlases. A recent community perspective emphasized that large cell atlases require reusable models that expose stable biological abstractions rather than dataset-specific embeddings [21]. This motivates reference models that define a fixed coordinate system into which new datasets can be mapped and compared. NMF is well suited to this role because it explicitly separates gene programs from cellular composition and supports projection of new data via non-negative least squares, enabling reuse without refitting the reference model.

This distinction matters operationally. When the coordinate system itself changes from study to study, comparisons are forced to occur at the level of ad hoc marker genes, curated signatures, or repeated retraining, each of which introduces ambiguity and weakens reproducibility. A fixed reference factor space instead supports measurement-to-program mapping as a key analysis step: once the reference is established, new datasets can be embedded into the same program coordinates, summarized at the program level, and compared directly to prior work without re-estimating latent structure. In theory, this is appealing, but in practice, reference NMF models have historically been trained on data corpora that were too small to yield models satisfactorily generalizable to new datasets.

Fitting interpretable factor models at Human Cell Census-scale requires algorithmic and computational advances beyond standard toolchains. Census-scale matrices are extremely sparse, exceed common indexing limits in standard R sparse matrix data structures, and NMF requires efficient access to both the expression matrix and its transpose during alternating updates of the weights matrices. We recently developed performant compressed data structures with value- and index-compressed sparse column formats designed for single-cell data to address memory limitations to achieve efficient in-core traversal and 64-bit indexing at atlas-scale [22], effectively reducing memory footprint almost 10x over R sparse matrices without compromising NMF performance. Coupled with our sparsity-optimized alternating non-negative least squares solvers [23, 17], it is now feasible to learn interpretable factor spaces directly on tens of millions of cells, rather than on heavily subsampled data, and to do so in-core without data distribution.

Here, we present an interpretable reference embedding of the Human Cell Census derived from non-negative matrix factorization. We factorized the normalized healthy human Census expression matrix into gene weights and cell weights at a fixed rank of 200 and released the resulting factorization as a reusable resource on the CZ cellxgene Census Models platform. Gene weights define transcriptional programs that can be labeled through gene-set enrichment using established resources [24, 25], while cell weights organize annotated cell types into neighborhoods in a shared program space. We further demonstrate reuse by projecting an independent cystic fibrosis dataset into the fixed reference programs and comparing factor structure with a new NMF model trained on the same data. This work provides a transparent, scalable program space that complements deep embeddings by prioritizing interpretability, stability, and direct reuse for atlas-scale single-cell analysis.

## Results

### An interpretable atlas-scale reference embedding of the Human Cell Census

We derived an interpretable reference embedding of the Human Cell Census by applying non-negative matrix factorization (NMF) to the normalized gene-by-cell expression matrix 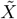. The factorization 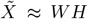 (rank *k* = 200) assigns a non-negative gene weight to each factor for every gene (columns of *W*) and a non-negative cell weight to each factor for every cell (columns of *H*), yielding an additive program representation of atlas-scale transcriptomes (Figure 1). This decomposition produces a shared coordinate system in which gene programs can be interpreted directly from *W* and cellular states can be interpreted directly from *H* based on simple inspection of the weights, without supervised labels or marker-based constraints during model fitting.

**Fig. 1:**
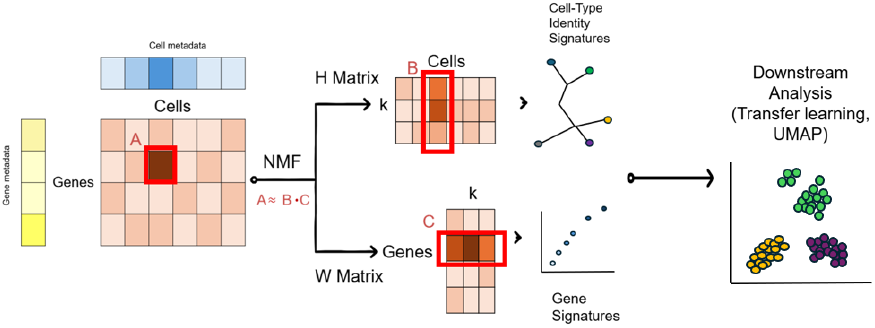
Non-negative matrix factorization yields interpretable gene and cell embeddings from single-cell expression data. A non-negative gene-by-cell expression matrix *X* is factorized as *X* ≈ *WH*, where *W* contains gene weights (genes × factors) and *H* contains cell weights (factors × cells). (A) Each observed entry of *X* is approximated by the dot product of a gene’s factor loadings in *W* (C) and a cell’s factor activations in *H* (B). The *H* matrix summarizes factor activity across cells and can be aggregated using cell-level metadata to derive cell-type signatures, while the *W* matrix captures gene-level contributions that define interpretable gene programs. Together, these matrices provide a low-dimensional, biologically interpretable latent representation that supports downstream analysis of cellular identity and transcriptional structure.

The resulting embedding separates gene-level and cell-level structure within a single model. Gene weights quantify the contribution of each gene to each factor, enabling program interpretation by gene ranking and gene-set enrichment. Cell weights quantify the contribution of each factor within each cell, enabling post hoc summarization across annotations and visualization in low-dimensional embeddings. This representation is compact, inspectable, and explicitly additive, supporting factor-wise reasoning about atlas-scale variation.

### Cell weights organize cell types into biologically relevant neighborhoods

We evaluated the structure of the cell weight matrix *H* by relating factor weights to standardized cell type annotations provided in the Cell Census. Mean cell weights aggregated by cell type revealed strong heterogeneity across factors, with many factors concentrating weight in restricted sets of biologically related annotations (Figure 2A–C). The observed concentration patterns indicate that the atlas-scale factors capture reproducible cellular programs. Specificity was apparent both in raw mean weights and in within-cell-type normalized weights, separating absolute magnitude from cell-type concentration.

**Fig. 2:**
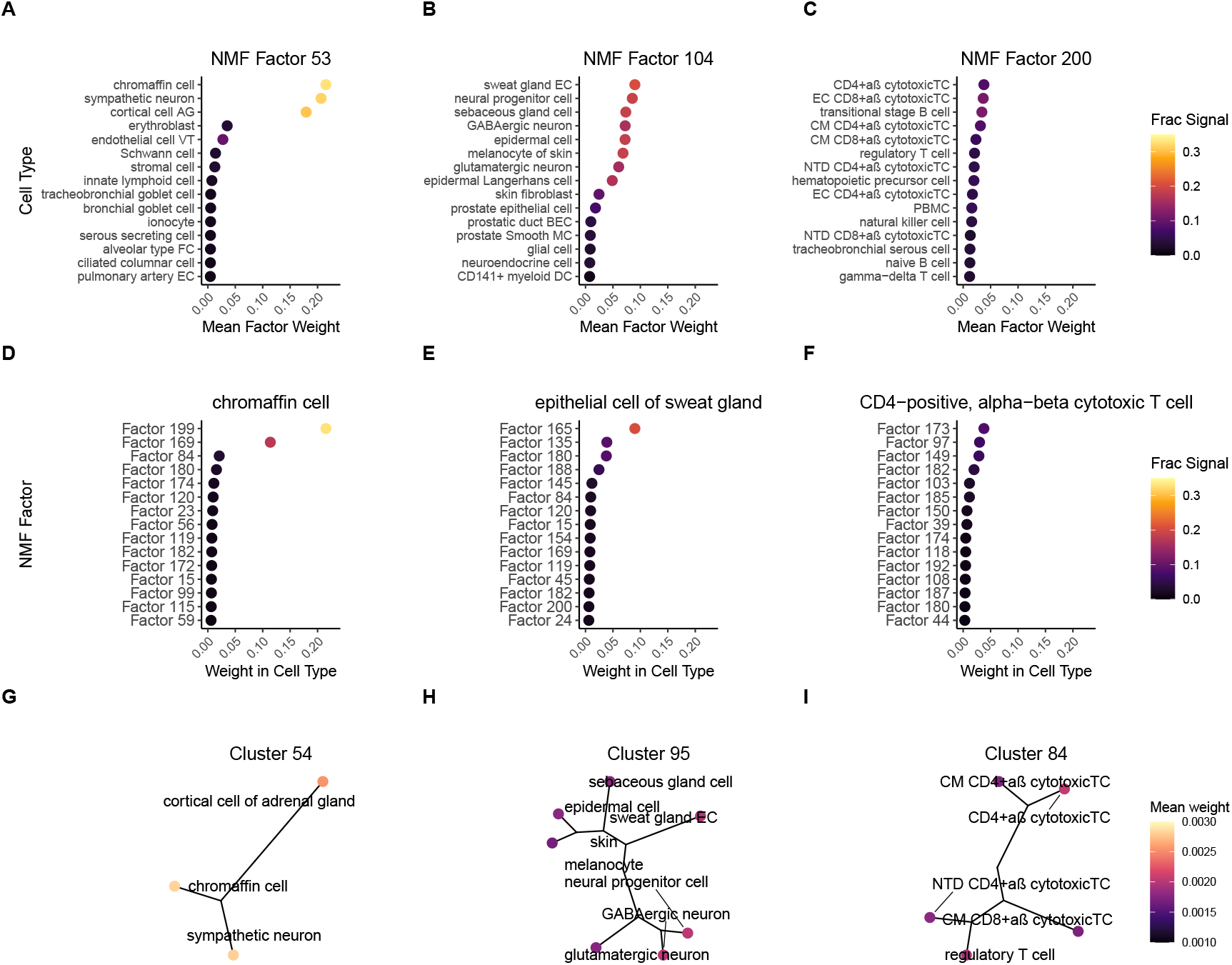
Cell-type organization revealed by the NMF cell weights matrix (*H*) across 28.5 million healthy human cells. (A–C) Factor-centric summaries for three representative factors, showing the mean weight assigned to each annotated cell type. Point size reflects mean factor weight, and color encodes the fraction of total factor signal contributed by each cell type, highlighting both specificity and reuse of transcriptional programs. (D–F) Cell-type–centric views for three representative cell types, illustrating the distribution of weights across factors and demonstrating that cell identities arise from combinations of multiple latent programs. (G–I) Unrooted trees derived from hierarchical clustering of cell types based on their *H* profiles, shown for three example clusters. Tip color indicates mean factor weight, revealing biologically coherent neighborhoods structured by shared and specialized transcriptional programs.

A complementary cell-type–centric view demonstrated that cell types are composed by distinct combinations of factors rather than by a single dominant program (Figure 2D–F). Some annotations were characterized by a small number of high-weight factors, whereas others distributed weight across many factors, consistent with differences in transcriptional complexity and heterogeneity. High-specificity factors were not necessarily those with the largest raw weights, motivating explicit reporting of both magnitude and within-type normalized weights in downstream summaries.

Cell-type relationships in program space were summarized by clustering cell types using their mean-weight profiles. Hierarchical clustering in factor-weight space grouped related lineages into neighborhoods that recapitulate biological structure (Figure 2G– I), including proximity between chromaffin-associated programs and sympathetic neuronal programs, and neighborhood structure among secretory epithelial populations. These results show that *H* provides an interpretable atlas-scale organization of annotated cell types in a common factor coordinate system.

### Gene weights define modular transcriptional programs with focused functional annotation

We assessed whether factor gene weights correspond to organized biological programs by analyzing columns of *W* with gene-set enrichment. For each factor, genes were ordered by their non-negative weights. Enrichment of this ordered rank vector against Gene Ontology gene sets showed many factors with distinct, sparse, and specific sets of genes that were enriched, indicating that many factors described different biological processes (Figure 3A). The global view of the GSEA results showed a number of factors that were non-specific or lacking enrichments, often a result of factors isolating batch signals or technical noise [26, 11]. Focused visualization of differentially enriched gene sets across representative factors highlighted interpretable programs spanning cilium organization and motility, extracellular matrix organization, mitochondrial function, immune activation, cell-cycle regulation, and nervous system development (Figure 3B). The inferred functional labels aligned with the cell-type structure observed in *H* (Figure 2), linking factor-associated cell-type neighborhoods to factor-associated gene programs.

**Fig. 3:**
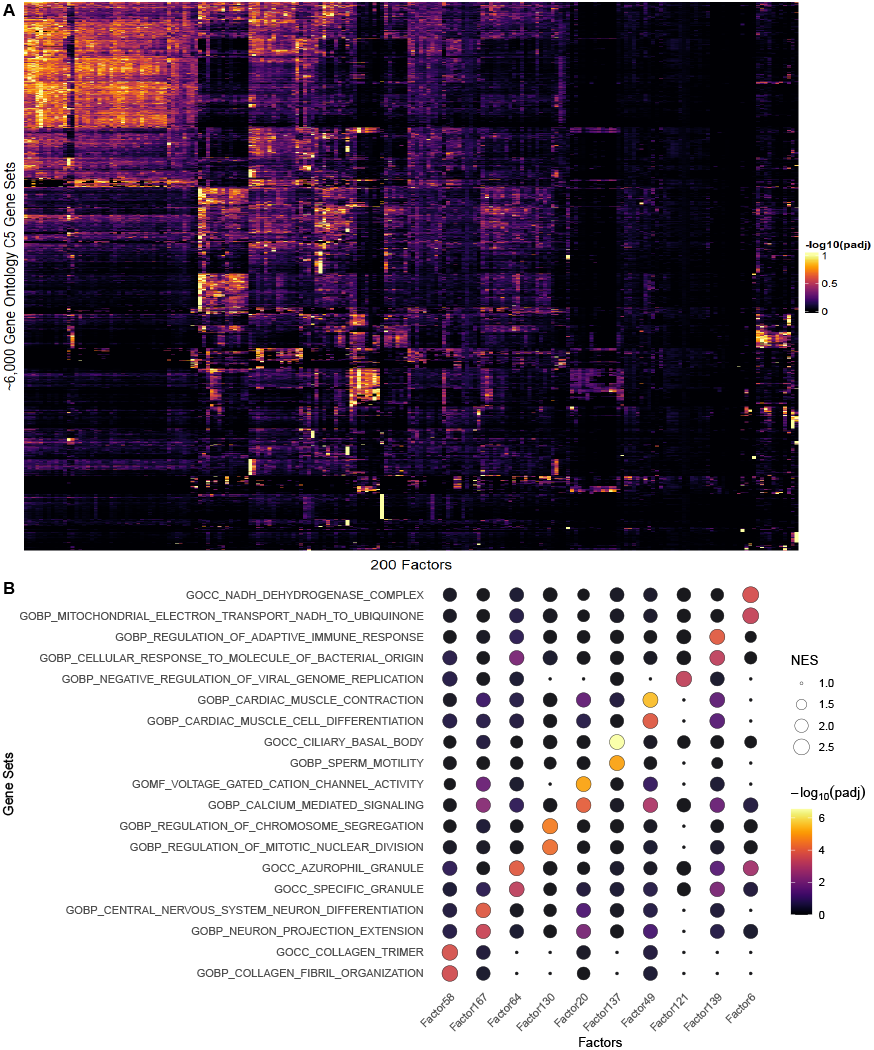
Biological programs identified by the NMF gene weights matrix (*W*). (A) Global landscape of pathway enrichment across all 200 factors, showing scaled − log_10_(padj) values for approximately 6,000 Gene Ontology (C5) gene sets. Each column corresponds to a factor and each row to a gene set, revealing that most factors concentrate signal into a small number of coherent biological pathways. (B) Focused view of ten representative factors highlighting their dominant gene-set associations. Bubble size indicates normalized enrichment score (NES), and color encodes enrichment significance, demonstrating that individual factors correspond to modular, biologically interpretable gene programs suitable for concise annotation.

These results establish that the gene weight matrix *W* supports direct and scalable program interpretation through gene ranking and enrichment while remaining mathematically consistent with cell-level organization in *H*, enabling factor-wise hypotheses grounded simultaneously in genes and annotated cellular structure.

### Reference projection preserves factor structure in an independent cystic fibrosis dataset

We tested reuse of the Census-derived reference factor space that was trained exclusively on healthy cells in an independent cystic fibrosis (CF) single-cell dataset of nearly 100,000 cells [27]. To do this, we projected new cells into the fixed reference (*W*_ref_) and compared the resulting cell weights to a new, independent NMF model trained *de novo* only on the CF dataset at the revealable rank (*k* = 70) determined by cross-validation. The new CF cell weights were obtained by solving a non-negative least squares problem for each cell using the fixed *W*_ref_, enabling direct comparison of reference-derived factor weights to independently learned weights *W*_*new*_ without modifying the reference programs (Figure 4A). Calculating the projection from *W*_*ref*_ took only about a minute on a standard CPU HPC node, while computing *W*_*new*_ required several hours on the same node to perform cross-validation and fit the final model, emphasizing the usability and convenience of leveraging our model as a reference.

**Fig. 4:**
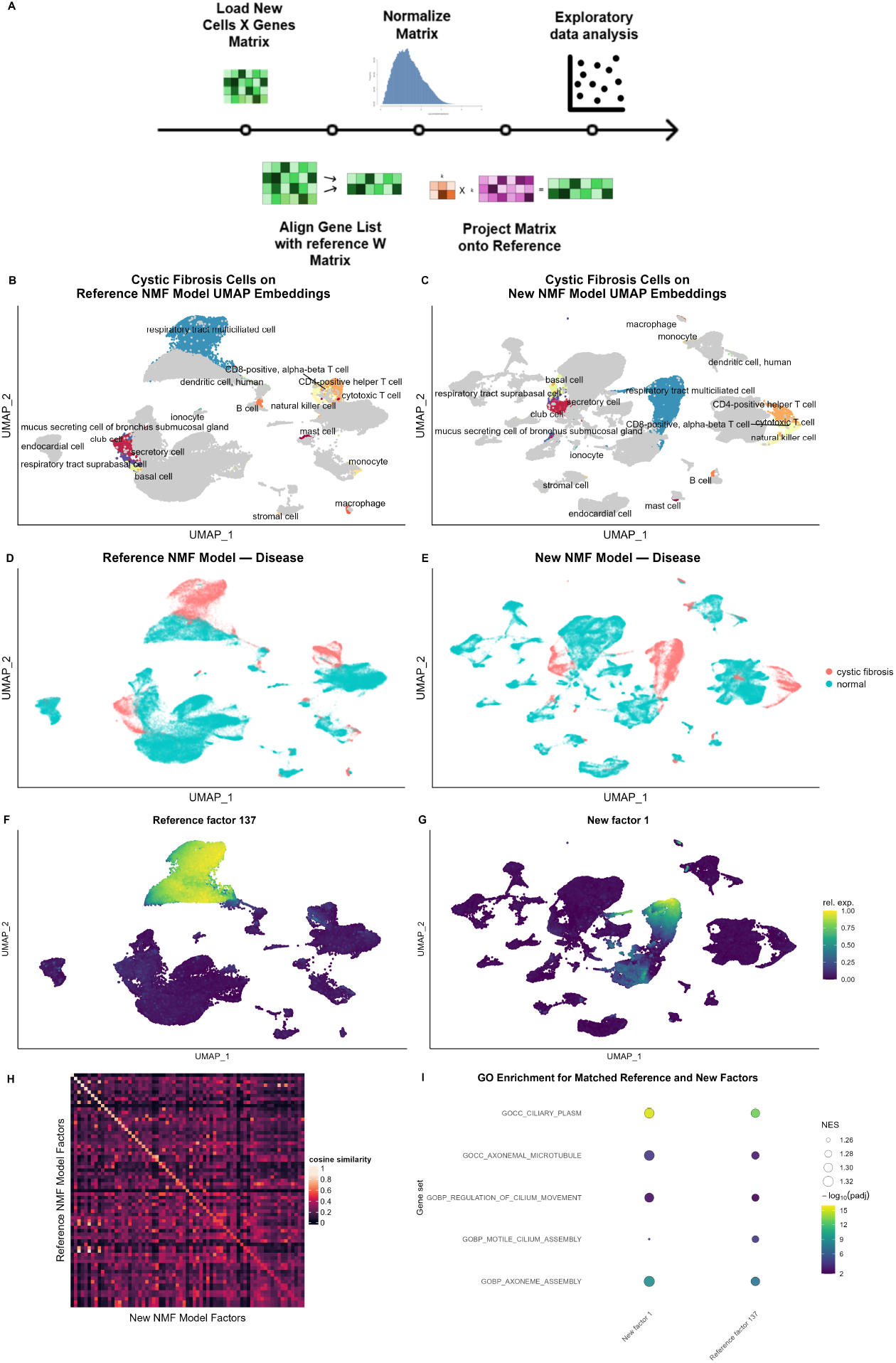
Reference projection embeds cystic fibrosis data into healthy Census NMF programs and recapitulates *de novo* structure. (A) Projection workflow for a new dataset: expression values are log-normalized, genes are aligned to the reference feature space, and per-cell factor weights are estimated by non-negative least squares against fixed reference gene programs *W*_ref_. In parallel, a *de novo* NMF model is fit to the same dataset at *k* = 70, selected by cross-validation, yielding (*W*_new_, *H*_new_). (B–C) UMAP embeddings computed from (B) projected factor weights *H*_proj_ and (C) *de novo* factor weights *H*_new_, with selected cystic fibrosis–associated cell types highlighted. (D–E) Disease status (CF vs. control) overlaid on the corresponding UMAPs. (F–G) Example matched transcriptional program visualized as scaled per-cell weights: (F) reference factor 137 and (G) *de novo* factor 1, both corresponding to a multiciliated, cilium-associated program. (H) Cosine similarity between gene-weight vectors in *W*_ref_ and *W*_new_, reordered by one-to-one bipartite matching and restricted to the *k* = 70 *de novo* factors. (I) Gene-set enrichment concordance for the matched factor pair in (F–G), showing consistent enrichment of cilium-related Gene Ontology terms.

Low-dimensional embeddings computed from factor cell weights using UMAP exhibited similar organization of annotated CF cell types under the reference projection and the new, independent model (Figure 4B–C). CF-associated populations, including respiratory tract multiciliated cells, epithelial subsets, and immune cell types, localized to comparable regions of the embedding under both representations and appeared equally distinct. Disease labels overlaid on the same embeddings revealed similar CF and normal cell distributions across the two models (Figure 4D–E), indicating that disease-associated structure emerges under projection despite the absence of CF samples during reference model training. Together, these results show that the reference programs trained on healthy data only can fit to a disease organization comparable to a model fit directly to the CF dataset, in this case, without any additional fine-tuning or retraining.

Factor correspondence between *W*_*ref*_ and *W*_*new*_ was quantified by cosine similarity between gene weight vectors, followed by one-to-one bipartite matching. The resulting similarity matrix exhibited a strong near-diagonal structure after reordering by the optimal assignment (Figure 4H), consistent with broad factor-wise correspondence between the two solutions and supporting NMF as a method for robustly learning convergent signatures across diverse datasets. A high-confidence matched pair (*W*_*ref*_ factor 137 and *W*_*new*_ factor 1) showed concordant localization in embedding space (Figure 4F–G) and was strongly enriched within the respiratory tract multiciliated cell population, including CF cells, supporting agreement at the level of factor-specific cellular structure.

Functional agreement between matched factors was further evaluated by gene-set enrichment analysis using the corresponding gene weights. The matched factor pair exhibited shared enrichment for cilium-related pathways with comparable enrichment direction and strength (Figure 4I), reinforcing correspondence at the level of interpretable biological programs. Together, these analyses demonstrate that a reference NMF model learned from healthy cells alone can be reused out of the box to organize, interpret, and annotate other datasets—even disease datasets— without retraining, while preserving factor identity, cell-type structure, and functional interpretability.

## Discussion

This study presents an interpretable, atlas-scale reference embedding of the Human Cell Census based on non-negative matrix factorization. By decomposing normalized gene-by-cell expression into non-negative gene weights and cell weights, the resulting model provides a shared coordinate system in which transcriptional programs can be inspected, summarized, and reused without retraining. The released factorization supports direct reasoning about both genes and cells at Census-scale, enabling exploratory analysis that remains transparent despite the size and heterogeneity of the underlying data.

Interpretability plays a central role in the utility of this representation. Additive factor structure enables explicit attribution of transcriptional signal to a limited set of programs, which can be aligned with Cell Census-provided metadata and functional annotations. The results show that factors concentrate signal into organized cell-type neighborhoods and focused gene programs rather than diffuse variation, even at tens of millions of cells. This structure supports program-level reasoning that is difficult to achieve with embeddings whose latent dimensions lack stable biological meaning or direct correspondence across datasets. The ability to embed new datasets without refitting preserves factor identity, reduces computational cost, and enables consistent comparison across studies and may be particularly beneficial for small or underpowered datasets. In the cystic fibrosis application, projected cell weights recapitulated the organization observed in an independent model, and matched factors exhibited concordant functional annotation. This demonstrates that the healthy reference programs capture stable–and even disease-relevant–transcriptional structure that generalizes beyond the data used for training, supporting their use as reusable analytical infrastructure.

Achieving this representation required methodological advances in scalable sparse computation and data structures that extend beyond standard single-cell toolchains. Conventional sparse matrix implementations encounter practical barriers at Census-scale due to indexing limits, memory overhead, and unfavorable access patterns, and alternating-update factorizations require efficient traversal of both *X* and *X*^*T*^. By enabling in-core optimization with compressed sparse formats and sparsity-aware alternating non-negative least squares updates, our implementation makes it feasible to learn an interpretable factor space directly on tens of millions of cells without subsampling or out-of-core operations. More generally, this illustrates how algorithmic and software design choices shape the scientific questions that are computationally accessible as single-cell atlases continue to grow.

Our approach has clear scope boundaries. The model is not designed to perform batch correction, integrate heterogeneous technologies, or optimize predictive performance relative to deep generative models, although batch factors are often isolated by NMF. At Census-scale, our NMF model will encode a library of batch and nuisance effects that will actually facilitate reference mapping where similar technical or assay effects are present. The rank in our model is fixed to provide a stable and tractable reference. Interpretability is emphasized over flexibility, and projection assumes a harmonized gene universe. These tradeoffs are deliberate and align with the goal of providing a transparent, reusable coordinate system rather than a universal integration solution.

These design choices also define predictable failure modes. Projection can only express new data as non-negative combinations of the reference programs, so truly novel cell states, rare disease-specific programs, or strong assay-specific artifacts that were not well-represented in the training data may be under-represented unless the reference is expanded or refit. In practice, this suggests a simple workflow: begin with projection for rapid, standardized program-level interpretation, and treat large residual structure, poorly explained cell groups, or consistently unassigned signals as evidence that a complementary new factorization (or an updated reference) is warranted. However, because NMF learns additive biological programs and has been trained on a large number of datasets, batch effects, sources of nuisance variation, and over 400 cell types across all major tissues, it is likely that the transfer of our reference embeddings will be sufficient in most cases.

An additional perspective is to treat rank as a tunable resolution parameter for biological abstraction. Lower ranks tend to emphasize coarse lineage structure and broadly shared programs, whereas higher ranks can separate finer activity states and more specific modules at the cost of increasing redundancy and susceptibility to overfitting in smaller datasets [28]. We fixed *k* = 200 to provide a stable reference that is expressive enough to capture diverse tissue programs while remaining tractable and inspectable. If programs in our reference are not applicable to new data, the projection simply sets weights mapping to those factors as 0. Future work could explore regularizations and methods that facilitate embeddings transfer optimized for interpretability and generalizability.

More broadly, this work underscores the need for scalable, interpretable models as single-cell atlases continue to grow. Linear, additive representations remain valuable complements to deep learning approaches, particularly when stable program definitions, cross-study comparability, and human interpretability are priorities. Reference embeddings such as the one presented here can support hypothesis generation, methodological benchmarking, and community-driven annotation, and may serve as scaffolding for future multi-resolution or multimodal extensions. As atlas-scale resources expand, combining scalable interpretability with expressive modeling will be essential for maintaining scientific insight alongside computational power [21].

## Methods

### Data acquisition and preprocessing

Gene-by-cell expression matrices and associated metadata were retrieved from the CZ cellxgene Discover portal (Human Cell Census; human-only; long-term supported release `2023-12-15`). Cells were filtered to include only primary data (`is_primary_data = TRUE`) from healthy donors (`disease = “normal”`) profiled using 10x Genomics 3^*′*^ or 5^*′*^ assays (v1, v2, or v3) or microwell-seq, with no tissue or cell-type restrictions. This yielded 28.5 million cells spanning all tissues and cell types in the Census. All genes in the Census feature space (approximately 60,700 genes) were retained. Raw counts were normalized per cell over total counts, multiplied by a scaling factor of 10^4^, and transformed with log1p. All downstream NMF model fitting, projection, and analyses were performed on this normalized representation.

### Memory- and compute-efficient sparse matrix data structures

NMF requires repeated access to both the expression matrix *X* and its transpose *X*^*T*^, which is infeasible at Census-scale using conventional sparse formats due to memory overhead and indexing limits. Representing *X* and *X*^*T*^ using `dgCMatrix` would require more than 10 TB of memory at Census-scale.

To enable fully in-core computation, we developed the Index- and Value-Compressed Sparse Column (IVCSC) format [22], optimized for zero-inflated, discrete-valued sparse matrices typical of single-cell transcriptomics.

The data-handling workflow when inputting blocked expression data handled within `singlet::run_nmf` involves a distributed transpose of *X* to yield *X*^*T*^, destructive conversion of both *X* and *X*^*T*^ to IVCSC format on a C++ backend, followed by the traversal of sparse columns in parallel without decompression during optimization.

IVCSC groups identical non-zero values within each column, stores each value once per group, and encodes each group of row indices using ordered, delta-encoded, byte-packed runs. This representation reduced memory usage by nearly tenfold relative to `dgCMatrix` while preserving efficient sparse–dense operations and column-wise OpenMP parallelization [22].

### Census-scale non-negative matrix factorization

We modeled the normalized expression matrix 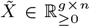 using non-negative matrix factorization (NMF),

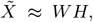

where 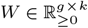 encodes gene-level transcriptional programs and 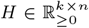 encodes cell-level program weights. The factorization rank was fixed at *k* = 200 to define a stable reference embedding.

To enable Census-scale fitting on tens of millions of cells, we developed a sparse-aware NMF implementation wrapped within the `singlet::run_nmf` function. In this function, normalized gene expression data from multiple `R dgCMatrix` objects were assembled into a single IVCSC matrix, after which the gene weight matrix *W* was initialized with random non-negative values. Model fitting proceeded via alternating non-negative least squares (NNLS) updates, first solving for the cell weight matrix *H* and then updating *W*, with an optional soft *𝓁*_1_ penalty (set to 0.01 in this study) applied to both matrices to encourage sparsity. Iterative updates continued until convergence, defined by a relative change in a correlation-based tolerance metric of tol ≤ 10^−4^ between successive updates of *W*.

Optimization converged in approximately 32 iterations for our model, with tolerance indicating a high-quality solution. Final model fitting was performed in-core without distribution on a high-performance CPU node (128 Intel Xeon threads, 1.5 TB RAM) over approximately one week of wall-clock time with RAM and thread utilization near capacity at all times. All operations were achieved in-core without distribution. In contrast, projecting new datasets into the learned reference space requires only seconds to minutes for hundreds of thousands of cells.

Exhaustive rank selection via cross-validation was infeasible at this scale. Instead, pilot analyses on representative subsets were used to identify a rank at this scale that would balance biological resolution, interpretability, and computational tractability, yielding the final choice of *k* = 200.

### Analysis of cell weights with cell metadata

Cell-level structure was analyzed by combining the factor weight matrix *H* with Census-provided cell-type annotations, which were not used during model fitting. For each annotated cell type, we computed mean factor weights across constituent cells and examined factor-centric summaries by ranking cell types according to their mean weight for each factor, as well as cell-type–centric summaries by ranking factors within each cell type. To distinguish factor specificity from absolute magnitude, mean factor weights were further normalized within each cell type by dividing by the total factor weight for that cell type. Relationships among cell types in program space were then assessed by agglomerative clustering based on their mean factor-weight profiles and visualized using unrooted trees and normalized factor contributions. This workflow enables interpretation of how transcriptional programs combine to form annotated cellular identities and how related cell types organize within the shared factor space.

### Analysis of gene weights with gene set enrichment

Gene-level transcriptional programs were interpreted by analyzing columns of the gene-weight matrix *W*. We began by subsetting the gene pathway universe to features present in the NMF model, and vice versa. Then for each factor, we performed the following: remove genes with zero weight, rank genes by non-negative weight, perform GSEA against Gene Ontology (C5) gene sets using `fgsea`, and record normalized enrichment scores (NES) and Benjamini– Hochberg adjusted *p* values (padj). These results were used to assign functional labels to factors and to assess concordance between gene-level programs and cell-level organization.

### Reference projection

To assess transferability of learned programs, we analyzed an independent cystic fibrosis (CF) dataset. Projection into the reference embedding proceeded as:

1. Normalize the new dataset using the same log-normalization procedure applied to the Census (it is equivalent to `Seurat::LogNormalize` with default parameters).
2. Align features (subset if necessary) in the new dataset and reference gene weights.
3. Use `singlet::ProjectData` to solve a non-negative least squares problem independently for each cell,

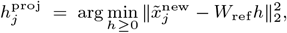

yielding projected cell weights directly comparable to the reference factors.

In parallel, a new NMF model was fit independently to the CF dataset to obtain (*W*_new_, *H*_new_) for direct comparison. We used `singlet::cross_validate nmf` to determine the optimal rank, defined as the rank achieving the lowest test set reconstruction error at convergence.

### Factor matching and embedding comparison

To assess factor correspondence between *W*_ref_ and *W*_new_, we paired factors using cosine similarity and bipartite matching. Matched factor pairs were used to compare localization in embedding space and concordance of pathway enrichment. Low-dimensional embeddings were computed from factor weights using UMAP (`uwot`) with cosine distance and `min_dist=0.3`. Disease status and selected cell-type labels were overlaid for visualization without influencing model fitting.

### Reproducibility and software

All analyses were performed using the open-source `singlet` framework and accompanying scripts. Random seeds were used to control initialization and optimization where possible. Exact numerical reproducibility of atlas-scale model fitting is not guaranteed due to non-determinism from parallel execution and floating-point arithmetic. However, factor structure and biological interpretations were robust across random initializations and closely related datasets.

## Software and data availability

This paper analyzes existing, publicly available single-cell RNA-seq data from the Chan Zuckerberg Initiative cellxgene Census (Human Cell Census), accessible at https://cellxgene.cziscience.com/census. All datasets used to train our reference NMF model use Census version 2023-12-15.

The Census-scale non-negative matrix factorization (NMF) reference embeddings reported in this study are publicly available through the CZ cellxgene Census Models portal (https://cellxgene.cziscience.com/census-models) as the nmf embedding for *Homo sapiens* (obs embedding: 28.5 million cells; var embedding: 60,700 genes). These embeddings are accessible programmatically via the Census API using `obs_embeddings=[“nmf”] and var_embeddings=[“nmf”]` for *H* and *W*, respectively.

The independent cystic fibrosis dataset used for projection and independent comparison (Figure 4) is publicly available via CZ cellxgene Discover as an AnnData `.h5ad` file at https://datasets.cellxgene.cziscience.com/170408c8-d0ec-441a-a1c1-261f7117763a.h5ad. This permanent download link references the dataset version used in this study (“Cystic Fibrosis and healthy control biopsies,” 96,479 cells; 10x 3’ v2/v3; *Homo sapiens*) associated with Berg *et al*. [27].

The non-negative matrix factorization and projection implementation used in this study is available in the open-source `singlet` package at https://github.com/zdebruine/singlet, in particular through the function `run_nmf` and `ProjectData`.

Code to reproduce all figures and analyses reported in this manuscript on the reference embedding, including projection, factor correspondence, and enrichment analyses applied to the released Census NMF embeddings, is available at https://github.com/liuyu-985/CellCensusNMF.

## Author contributions statement

Conceptualization, Z.J.D. and T.J.T.; methodology, Z.J.D. and Y.-T.L.; software, Z.J.D. and Y.-T.L.; investigation, Y.-T.L. and Z.J.D.; data curation, Y.-T.L.; formal analysis, Y.-T.L.; visualization, Y.-T.L.; writing–original draft, Z.J.D.; writing– review & editing, Y.-T.L., T.J.T., and Z.J.D.; supervision, Z.J.D.; resources, Z.J.D.; funding acquisition, Z.J.D.; validation, Y.-T.L. and Z.J.D.; investigative support, T.J.T. All authors approved the final manuscript.

